# The disordered and structured regions of α-Synuclein contribute to membrane remodeling synergistically

**DOI:** 10.64898/2026.03.19.713051

**Authors:** David H. Johnson, Jia Shiuan Liow, Orianna H. Kou, Wade F. Zeno

**Affiliations:** Mork Family Department of Chemical Engineering and Materials Science, University of Southern California, Los Angeles, California 90089, United States; Department of Physics and Astronomy, University of Southern California, Los Angeles, California 90089, United States

**Author notes:** **Corresponding Author** Wade F. Zeno.

## Abstract

α-Synuclein (αSyn) remodels cellular membranes through interactions that involve both its structured, membrane-binding N-terminal domain (NTD) and intrinsically disordered C-terminal domain (CTD). While the amphipathic NTD helix is known to insert into lipid bilayers and generate curvature, the contribution of the acidic CTD remains unclear. Here, we dissect the individual and cooperative roles of these domains using Supported Bilayers with Excess Membrane Reservoir (SUPER) templates to quantify membrane remodeling via membrane fission and membrane morphological deformations (i.e., membrane budding and tubulation). We show that both the NTD and CTD independently remodel membranes, while full-length αSyn exhibits greater remodeling ability than either the NTD or CTD in isolation. This result demonstrates a synergistic amplification between helix insertion of the NTD and the tethered, disordered CTD. To further probe the mechanism of membrane remodeling by the CTD, we modulated the chain length of the protein, the bulk ionic strength of the solution (i.e., charge screening), and applied relevant polymer scaling laws for disordered proteins. Our results suggest that the membrane remodeling mechanism for the disordered CTD is electrostatic in nature, stemming from protein-protein repulsion at elevated binding densities. Together, our findings reveal a cooperative energetic mechanism in which N-terminal helix insertion biases membrane curvature and the disordered, C-terminal domain adds an additional electrostatic component that helps to overcome the free energy barrier for membrane bending.

## INTRODUCTION

α-Synuclein (αSyn) is an intrinsically disordered protein (IDP), enriched in the pre-synaptic terminal of neurons^1^, where it plays an important role in synaptic vesicle trafficking and membrane remodeling^2–5^. αSyn is also implicated in a variety of neurodegenerative disorders collectively termed synucleinopathies^6–7^, where, for example, it has been shown to pathologically remodel mitochondrial membranes ^8–9^. These membrane remodeling activities arise from αSyn’s interactions with lipid bilayers, yet the molecular features of the protein that drive these deformations remain incompletely understood ^10–12^. In particular, αSyn contains distinct structural regions: an amphipathic, membrane-interacting N-terminal domain (NTD) and an acidic C-terminal domain (CTD). While the full-length protein is disordered in solution, the NTD adopts an α-helical structure upon membrane association, whereas the CTD remains disordered and tethered to the membrane surface (Figure 1A). Determining how these domains individually and collectively contribute to membrane remodeling remains an important unresolved question.

**Figure 1.**
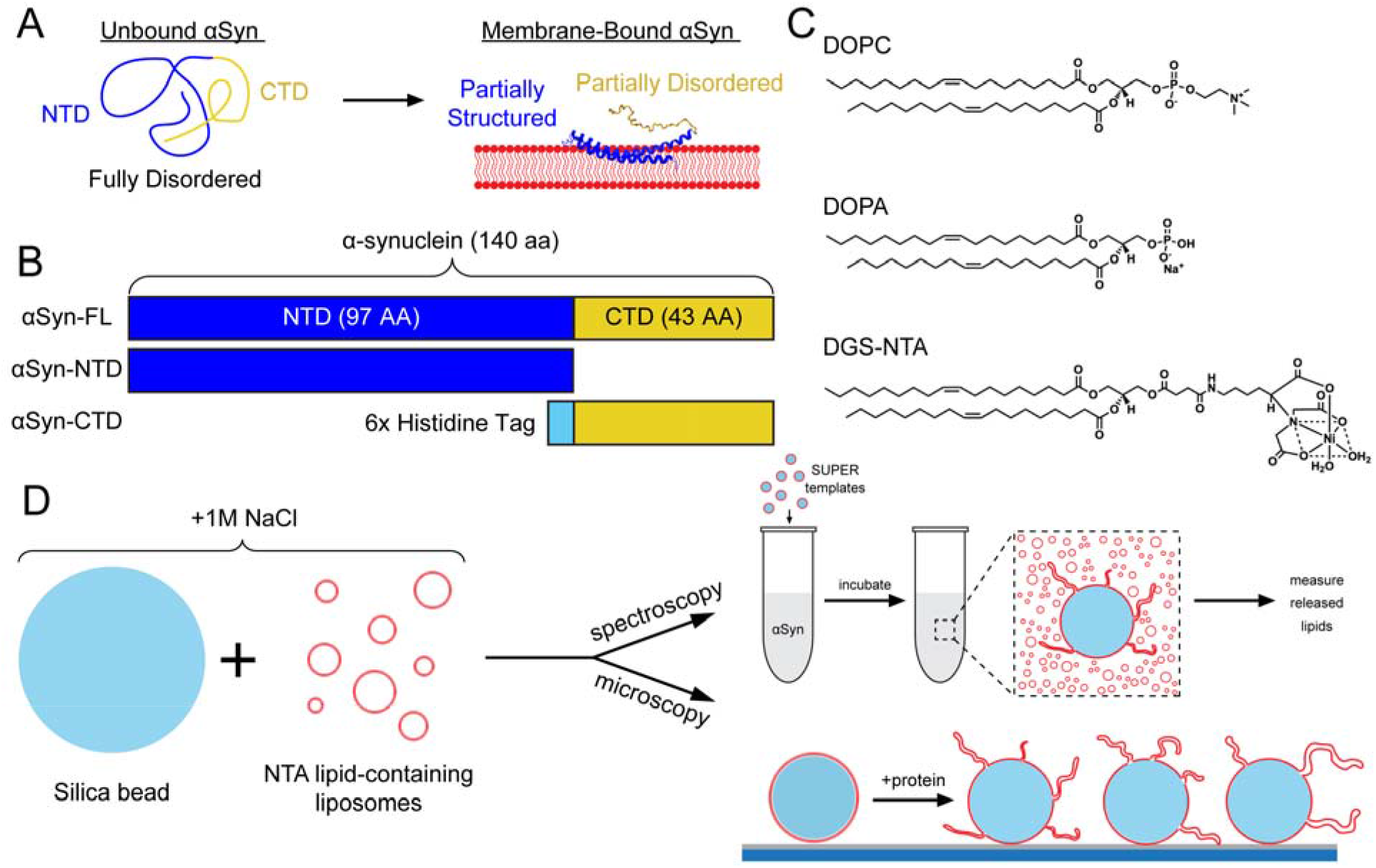
Schematics representing the experimental setup for the study. **(A)** Representative structure of αSyn in its unbound and membrane-bound states. The bound structure of αSyn was adapted from UniProt P61144. **(B)** The three αSyn-derived constructs used for membrane remodeling studies. **(C)** Chemical structures of the primary lipid components used in the SUPER template assays. **(D)** Schematic of SUPER template synthesis (left), spectroscopy-based membrane fission assay (top right), and microscopy-based membrane deformation assay (bottom right).

A large body of work has examined the molecular mechanisms by which αSyn deforms lipid membranes^10–16^. Mechanistic studies have largely focused on the membrane-bound NTD, using simulations, supported lipid bilayers, and electron microscopy to examine how αSyn perturbs bilayer structure ^12, 17–19^. These studies suggest that insertion and amphipathic interactions of the NTD can induce membrane thinning, lateral expansion of lipid headgroups, and curvature generation. Despite these advances, comparatively little attention has been given to the potential role of the disordered CTD in membrane remodeling ^18, 20–21^. As a result, the extent to which the two domains contribute individually to membrane deformation, and how their combined presence influences remodeling by the full-length protein, remains unclear.

Insight into possible contributions from the CTD can be drawn from studies of other membrane-associated IDPs. Membrane-anchored IDPs can generate curvature through steric pressure arising from their extended conformations. Because disordered chains occupy a larger effective volume than folded proteins of comparable molecular weight, crowding of IDPs at membrane surfaces can produce steric pressure sufficient to drive membrane bending ^22–24^. Consistent with this mechanism, shortening membrane-tethered disordered domains reduces their ability to deform membranes ^25^. Although many systems examined to date involved IDPs composed of several hundred residues^26–27^, the CTD of αSyn is much shorter at approximately 40 residues in length. Despite its modest length, the CTD is highly enriched in negatively charged aspartic and glutamic acid residues, giving the sequence a high net charge that could promote electrostatic repulsion between neighboring IDPs on the membrane surface. Such electrostatic interactions may augment steric crowding and enhance curvature generation. Additionally, the bending ability of disordered domains can be amplified when they flank a structured membrane-binding domain ^28–29^, as occurs for the CTD positioned adjacent to the N-terminal helix of αSyn. Whether these steric and electrostatic crowding mechanisms contribute to αSyn-mediated membrane remodeling remains unresolved.

In this study, we dissect the contributions of the structured NTD (αSyn-NTD) and the intrinsically disordered CTD (αSyn-CTD) to αSyn-mediated membrane remodeling using truncation mutants. Using Supported Bilayers with Excess Membrane Reservoir (SUPER) templates, we quantify membrane fission by fluorescence spectroscopy and characterize membrane morphology by confocal microscopy. We find that both domains independently induce membrane fission, while full-length αSyn exhibits enhanced remodeling, indicating synergistic coupling between helix insertion and disordered-domain-mediated pressure. By varying CTD length and ionic strength, we further show that electrostatic repulsion is a dominant driver of CTD-mediated remodeling. Together, these results establish a cooperative mechanism in which helix-mediated membrane binding and disordered electrostatic pressure act in concert to enhance membrane deformation by αSyn.

## EXPERIMENTAL MATERIALS AND METHODS

### Materials

1,2-dioleoyl-sn-glycero-3-phosphocholine (DOPC), 1,2-dioleoyl-sn-glycero-3-phosphate (sodium salt) (DOPA), and 1,2-dioleoyl-sn-glycero-3-[(N-(5-amino-1-carboxypentyl)iminodiacetic acid)succinyl] (ammonium salt) (DGS-NTA) were purchased from Avanti Research. Dipalmitoyl-decaethylene glycol-biotin (DP-EG15-biotin) was provided by D. Sasaki from Sandia National Laboratories, Livermore, CA^30^. ATTO 647□N 1,2-Dipalmitoyl-snglycero-3-phosphoethanolamine (ATTO647N-DPPE) and ATTO 488-maleimide were purchased from Leica Microsystems. Isopropyl-beta-D-thiogalactoside (IPTG) and Dithiothreitol (DTT) were purchased from Gold Biotechnology. Tris(2-carboxyethyl)phosphine hydrochloride (TCEP), phenylmethanesulfonyl fluoride (PMSF), sodium phosphate monobasic, and sodium phosphate dibasic were purchased from Sigma-Aldrich. 4-(2-hydroxyethyl)-1-piparazineethanesulphonic acid (HEPES), poly-L-lysine (PLL), and Tris(hydroxymethyl)aminomethane (Tris) were purchased from Fisher Scientific. 2.5 µm non-functionalized silica spheres were purchased from Bangs Labs. 9.2 µm monodisperse silica spheres were purchased from Cospheric. Amine-reactive PEG (mPEG-SVA, MW 5000) and PEG-biotin (Biotin-PEG-SVA, MW 5000) were purchased from Laysan Bio, Inc.

### Plasmids for Microscopy and Spectroscopy Assays

The wild-type α-synuclein (FL-αSyn-WT) plasmid (pET21a backbone) was provided by Peter Chung (University of Southern California, Los Angeles, CA). The Y136C full-length αSyn (FL-αSyn-Y136C) mutation (pRK172 backbone) was provided by Jennifer C. Lee (National Institutes of Health, Bethesda, MD). All other constructs: pET21a α-synuclein 1-97 (αSyn-NTD), pET21a α-synuclein 1-97 G93C, pGex4T2 his-α-synuclein 98-140 Y136C (his-αSynCTD), and pGex4T2 his-α-synuclein 98-140(x3) Y136C (his-αSynCTDx3) were synthesized by GenScript Biotech Corporation (Nanjing, China).

### Protein Expression and Purification

Wildtype and Y136C mutated αSyn-FL and αSyn-FL (sequences in Supplementary Table 1) were transformed into BL21 (DE3) cells. To produce the acetylated versions, the plasmids were transformed into BL21 (DE3) containing the pNatB plasmid coding for N-α-acetyltransferase to acetylate the N-terminus of the protein^31^. The purification was performed as described previously^32^. Briefly, cells were incubated in 2xTY media until an OD_600_ ~0.6. The cells were pelleted, resuspended, and sonicated. DNase I (Thermo Fisher Scientific) was added to the lysate, followed by acid precipitation by adjusting the pH to 3.5. The cell debris was pelleted, and a 50% ammonium sulfate precipitation was performed, followed by another pelleting. The pellet was resuspended in 20 mM Tris (pH 8.0), loaded onto a 5 mL HiTrap Q FF column (Cytiva), and eluted with 1 M NaCl. The eluted protein was concentrated using a 3 kDa MWCO filter (Sigma-Aldrich) and run through SEC on a Superdex 200 10/300 column (Cytiva). The final fractions were concentrated and snap frozen in LN2. The extinction coefficients were 5960□M^−1^□cm^−1^ and 4470□M^−1^ cm^−1^ for FL-αSyn-WT and FL-αSyn-Y136C, respectively.

αSyn-NTD (either with or without a G93C mutation, see sequences in Supplementary Table 1) were transformed into BL21 and pNatB-BL21 (DE3) as described above. After incubation in 2xTY at 37 °C to an OD600 of ~0.6, protein production was induced with 1 mM IPTG for 5 hours. Cells were pelleted, then resuspended in 25 mM Tris, 20 mM NaCl, 1 mM EDTA (pH 8) with 1 mM PMSF and 1x Roche complete protease inhibitor tablet (Sigma-Aldrich). 5 mM DTT was added to the G93C mutant to prevent disulfide bond formation. The cells were sonicated on ice at 65% amplitude for 16 minutes. 10X DNase reaction buffer (100 mM TRIS, 25 mM MgCl_2_, 1 mM CaCl_2_, pH 8) was added to the sonicated cells along with DNase I, and the mixture was incubated at 37 °C for 30 minutes. 50 mM EDTA was added to stop the DNase reaction, and the solution was incubated at 65 °C for 10 minutes. Acid precipitation was performed by adjusting the solution pH to 3.5 with 5 M HCl added dropwise. For the αSyn-NTD, the solution was pelleted for 25 minutes at 20,000 x g. The supernatant was then concentrated to 2 mL using 3 kDa MWCO centrifugal filters. For the G93C construct, a two-step ammonium sulfate precipitation was performed after acid precipitation. 0.116 g/mL of ammonium sulfate was added to the solution at 4 °C, and the mixture was stirred for 1 hour. The solution was spun down for 25 minutes at 25,000 x g. 0.244 g/mL of ammonium sulfate was added to the supernatant, which was then spun at 4 °C for 1 more hour. The centrifugation was repeated, and the pellet was resuspended in 1 mL of 25 mM HEPES, 0.5mM TCEP (pH 7.4). Both proteins were then run on SEC over a HiLoad 16/600 75 pg column in 25 mM HEPES and 2 M NaCl. Selected fractions were concentrated, diluted to 500 mM NaCl by adding 25 mM HEPES, and snap-frozen in liquid nitrogen. The extinction coefficient was 1490□M^−1^□ cm^−1^ for both NTD mutants.

αSyn-CTD and αSyn-CTDx3 (both containing a single cysteine mutation, see sequences in Supplementary Table 1) were expressed as fusion proteins containing N-terminal glutathione-S-transferase (GST) in BL21 (DE3) cells. After reaching OD_600_ ~ 0.6 at 37 °C in 2xTY, cells were induced by adding 100 µM IPTG and incubating at 37 °C for 3 hours. Cells were then pelleted and resuspended in 500 mM Tris, 5% v/v glycerol (pH 8), with 5 mM DTT, 1 mM PMSF, and 1x Roche Complete Protease Inhibitor Tablet. Triton-X was added to the solution to a final concentration of 1% v/v. The cells were sonicated at 60% amplitude for 16 minutes on ice then pelleted for 40 minutes at 103,000 x g. Proteins were then purified by incubation with glutathione resin beads (Thermo Fisher Scientific) for 1 hour at 4 °C, then placed into a column and washed consecutively with 400 mM Tris, 1 M NaCl (pH 7.4), 1 mM PMSF, 5 mM DTT, and 25 mM HEPES, 150 mM NaCl (pH 7.4), 0.5 mM TCEP. The protein was cleaved from the GST beads by incubation with thrombin (Sigma-Aldrich) at 4 °C overnight. Thrombin was removed by incubating p-aminobenzamidine-agarose beads (Sigma-Aldrich) in the solution for 1 hour at 4 °C. The proteins were concentrated with 3 kDa MWCO centrifugal filters and snap-frozen in liquid nitrogen. The extinction coefficients were 2980□M^−1^□cm^−1^ and 11920□M^−1^□cm^−1^ for αSyn-CTD and αSyn-CTDx3 mutants, respectively.

### Fluorescent Labeling

Fluorescent labeling of all cysteine-containing mutants was performed as described previously^32^. Briefly, ATTO-488 maleimide was added to the unlabeled proteins at a 4X molar excess of dye. Reactions were performed for 2 h at room temperature or overnight at 4 °C. Excess dye was removed after labeling by SEC on a Superdex 75 10/300 GL (Cytiva), and the labeling ratio was confirmed to be 1:1 for all reactions by UV-Vis spectroscopy.

### SUV Preparation

Lipid aliquots in chloroform were thawed from −80 °C and combined in a glass tube at the following molar percentage: 63% DOPC, 25% DOPA, 10% DGS-NTA, 1% DP-EG15-biotin, 1% ATTO 647-DPPE. The chloroform was evaporated with a nitrogen stream, then placed under vacuum for at least 1 h. The resulting lipid film was rehydrated with either MilliQ water for super template preparation or 20 mM sodium phosphate (pH 7.0), with the appropriate salt concentration for imaging (i.e., 10 mM, 150 mM, or 1 M NaCl) for the tethered vesicle assay. After 5 min of hydration, vesicles were extruded through a 100 nm polycarbonate membrane (Whatman), stored at 4 °C, and used within one week.

### SUPER Template Preparation

SUPER templates were prepared according to published protocols ^28, 33^. Using the 100 nm extruded SUVs rehydrated in MilliQ water, SUPER templates were prepared by combining 50 μL of 200 μM SUVs, 50 μL of 2 M NaCl, and 5 × 10^6^ silica beads in a low-bind centrifuge tube. The mixture was gently stirred for 30-45 minutes at 4 °C. Excess vesicles were removed by four rounds of centrifugation (300 × g, 2 min) and resuspension in 1 mL MilliQ water. The final templates were stored on ice for up to 3 hours before use.

### Super Template Fission Spectroscopy Measurement

Membrane fission from SUPER Templates was quantified as described previously ^28^. Briefly, 45 µL solutions of 20 mM sodium phosphate (pH 7.0) with the appropriate salt and protein concentration with 5 mM TCEP were created in low-adhesion microcentrifuge tubes. 5 µL of SUPER templates were pipetted to the top of the solution and allowed to settle for 30 minutes with no agitation. The tubes were then gently centrifuged (2 mins, 300 x g) to separate SUPER templates with intact membranes from the released lipids in solution. 37.6 µL of supernatant was transferred to a 96-well plate and mixed with buffer containing 0.1% Triton X-100 and 5 mM TCEP to reach a final volume of 100 uL in the well. The samples were measured on a Biotek Synergy Neo2 plate reader (Agilent) with a 620/680 polarizing cube. Three controls were implemented to calculate the total fission products from the templates.. The first control measured a blank solution of 0.1% Triton-X in the same buffer used in the experiments and was subtracted from all samples to eliminate background fluorescence due to the buffer. The second control measured the fluorescent signal from a fission experiment performed with the SUPER templates in the absence of protein to measure the released lipid due to template processing (centrifugation and pipetting), which was also subtracted from every sample. The first two controls are fluorescent controls that represent the lower bounds of the fluorescent fission measurements (i.e. when no protein is present to cause fission). The third control, serving as the upper bound of fission, measured the solubilized lipid from SUPER templates added directly to a solution of 0.1% Triton-X, solubilizing all lipid on the template. The percentage of fission products was calculated by dividing the fluorescence after protein incubation by the fluorescence of the third control.

### Slide Passivation and Membrane Tethering

Glass coverslips were passivated as previously described^34–35^. Briefly, glass coverslips were passivated with a layer of biotinylated, 5 kDa, PLL-PEG for all imaging experiments. For the tethered vesicle assay, NeutrAvidin was added to tether the SUVs to the glass surface.

PLL-PEG was synthesized by combining mPEG-SVA (98%) with Biotin-PEG-SVA (2%) to a 20 mg/mL mixture of PLL in 50mM sodium tetraborate (pH 8.5) for a lysine to PEG ratio of 5:1. The mixture was stirred at room temperature for 6 hours and buffer exchanged into 25mM HEPES, 150mM NaCl (pH 7.4) using a 5mL, 7 kDa MWCO Zeba Spin column (Thermo Fisher Scientific).

Imaging wells were made from 5 mm diameter holes in 0.8 mm thick silicone gaskets (Grace Bio-Labs). The coverslips were cleaned with a 2% v/v Hellmanex III (Hellma Analytics), rinsed with MilliQ water, and dried with a nitrogen stream. The gaskets were placed on the glass slide, and 30 μL of PLL-PEG was added to the wells, which were then incubated for 20 minutes. Excess PLL-PEG was rinsed using gentle pipetting with 20 mM sodium phosphate, 150 mM NaCl (pH 7.0) (Preparation Buffer).

For binding curves, extruded vesicles were tethered within imaging wells. Vesicles were tethered by first incubating the imaging wells with 6 µg of NeutrAvidin for 10 minutes. Excess NeutrAvidin was rinsed with Preparation Buffer. The prepared 100 nm extruded SUVs were then added to the wells at 1 μM and incubated for 10 minutes. Excess vesicles were rinsed with Preparation Buffer. The buffer composition was then exchanged to the imaging salt conditions (10 mM, 150 mM, or 1 M NaCl) with 5 mM TCEP. The proteins were then added to the wells at the appropriate concentrations for imaging and incubated for 10 minutes before imaging.

For membrane morphological deformation analysis, SUPER templates were allowed to settle within imaging wells and were not tethered by omitting NeutrAvidin from the well preparation. The PLL-PEG passivated slides with 30 μL of appropriate imaging buffer were incubated with 1 μL of the prepared 9.2 μm SUPER templates and allowed to incubate for 10 minutes. The wells were rinsed once with the same buffer to ensure they were equilibrated with the appropriate protein imaging buffer. Protein was then added at the appropriate concentrations and incubated for 15 minutes before imaging. Wells were imaged one at a time to ensure equivalent incubation times.

### Confocal Microscopy

All imaging wells were imaged on a Leica Stellaris 5 laser scanning confocal microscope as described previously^32^. Briefly, two excitation lasers were used: 488 nm and 647 nm for the αSyn and the SUVs, respectively. The detection wavelengths for each laser were 493 – 558 nm and 643 – 749 nm for the 488 and 647 lasers, respectively. A Leica HC PL APO 63x, 1.4 NA oil immersion objective was used to acquire the images. For tethered vesicle analysis, the zoom factor was set such that each image was composed of square 70×70 nm pixels and all images were acquired with a 400 Hz scan speed within a single confocal imaging plane near the glass cover slip. For SUPER template imaging, the following changes were made to the image acquisition settings: the scan speed was adjusted to 700 Hz, the zoom was set such that each pixel was a 283×283 nm square, and each field of view was a 25 image z-stack to capture all features on the SUPER template.

### Confocal Microscopy Image Processing

For the tethered vesicle assay, the image processing was performed exactly as previously described ^32, 34^. Images of diffraction limited puncta in the αSyn and SUV fluorescent channels were processed using publicly available particle detection software (CMEAnalysis) ^36^. All puncta over 250 nm were excluded from the analysis to ensure all puncta were diffraction-limited. Any puncta not significantly above the background noise were also excluded from the analysis. Protein binding was determined by analyzing colocalization between the two fluorescent channels.

For the SUPER template assay, z-stacks were analyzed frame by frame in ImageJ to determine the percentage of total templates with features, the number of features per template, and the type of feature on the template surface. A feature was defined as any protein and lipid colocalized structure protruding from the surface of the template. The features were sorted into 3 bins based on the observed protrusions: membrane buds, thick tubules, and sub-diffraction limited tubules. Membrane buds are defined as spherical structures attached to the template surface of any diameter. Sub-diffraction limited tubules are defined as tubules ≤ 300 nm in diameter and thick tubules are ≥ 300 nm in diameter. Two additional bins were created based on the length of the tubules. Long features had a total length > 2 µm while short features had a total length of ≤ 2 µm.

### Calibration of Vesicle Diameters

SUV diameters in the tethered vesicle assay were calibrated as described previously ^32, 34–35, 37^. Briefly, SUVs were imaged to obtain a distribution of fluorescence intensities for the vesicle population. The conversion factor between vesicle diameter and fluorescence intensity was determined by overlaying the square root of the fluorescence intensity with the DLS diameter distribution. Vesicles included in the binding analyses were selected from the overall size distribution to be 80-120 nm in size, resulting in an average size of 100 nm, to correct for curvature sensing effects of the protein in our results. All other vesicles were excluded from the analysis.

### Calibration of Number of Bound Proteins and Conversion to Surface Coverage

The number of proteins bound to vesicles of a particular size was determined as described previously ^32^. Briefly, each αSyn mutant used to image was adsorbed to a glass coverslip, and single-molecule imaging was performed. From the distribution of fluorescence intensities, the intensity of a single protein was determined from the peak of this distribution, representing the intensity of a single ATTO488-labeled protein.

The average number of proteins on 100 nm vesicles was plotted for each protein and concentration to create binding isotherms (Fig. S1). Each curve was fit to the following equation for a Langmuir Isotherm:

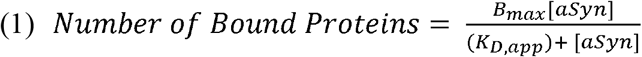

Where B_max_ is the maximum value of protein bound, [αSyn] is the bulk concentration of αSyn construct incubated with the vesicles, and K_*D,app*_ is the apparent dissociation constant. To convert the amount of binding on tethered vesicles to the relative binding in the SUPER template assays, the K_*D,app*_ can be scaled using the lipid concentrations in both assays to convert protein concentrations to relative bound densities of protein, as seen in Figs. 2B, 2D, S2C, S5B, and S5E.

**Figure 2.**
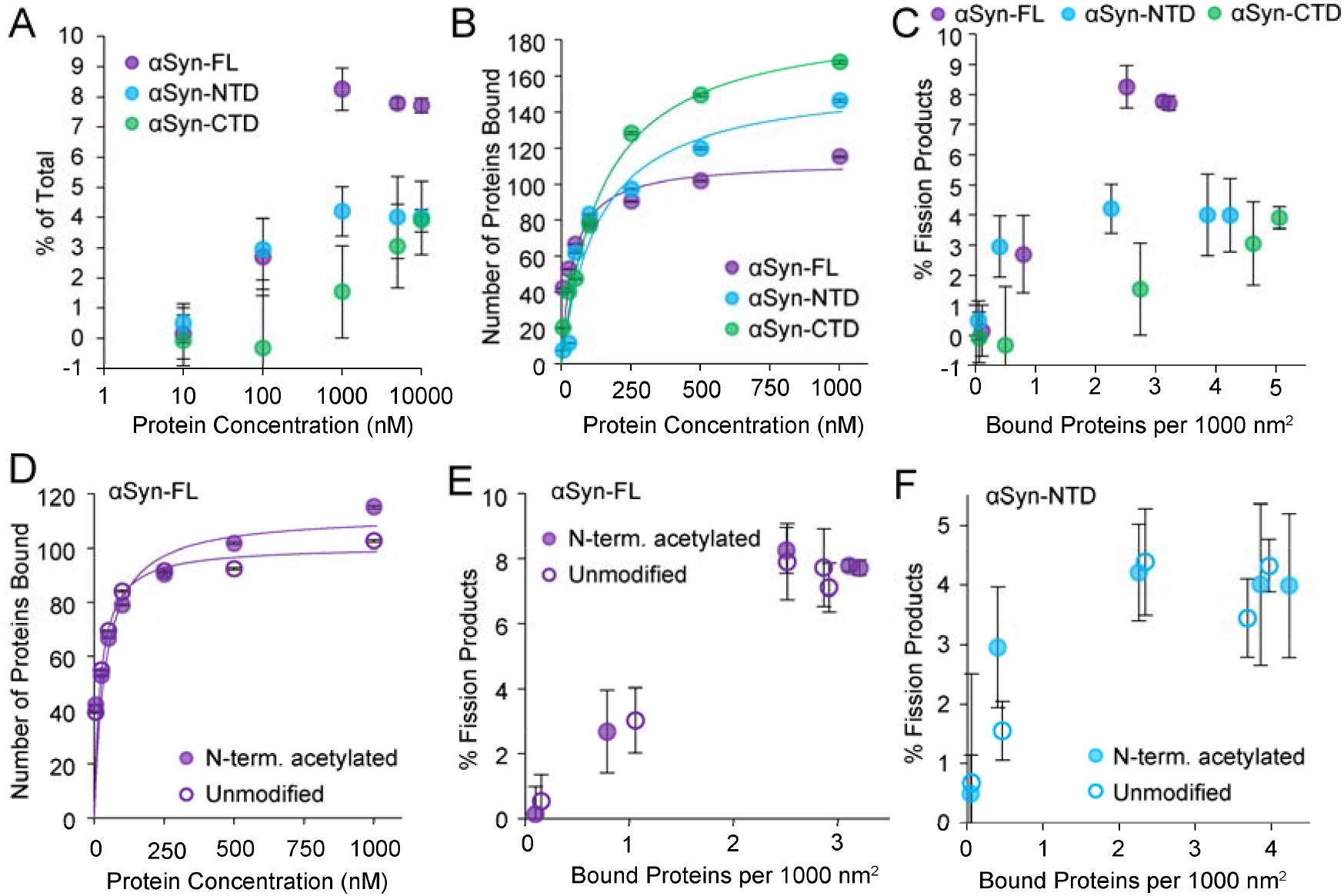
Membrane fission by αSyn truncation mutants and effect of N-terminal acetylation. **(A)** Percent of fission products as a function of protein concentration for αSyn-FL, αSyn-NTD, and αSyn-CTD. **(B)** Binding isotherms with Langmuir fits obtained from tethered vesicle assay. N = 2058-2169 imaged vesicles for αSyn-FL, N = 1579 – 2117 imaged vesicles for αSyn-NTD, and N = 886-996 imaged vesicles for αSyn-CTD. **(C)** Percent of fission products plotted versus equivalent surface coverage for each αSyn construct. **(D)** Binding isotherms with Langmuir fits obtained from tethered vesicle assay for acetylated and unmodified αSyn-FL. N = 2058-2169 imaged vesicles for acetylated FL and N = 1839 – 1920 imaged vesicles for unmodified FL. **(E)** Percent of fission products plotted versus equivalent surface coverage for acetylated and unmodified αSyn-FL. **(F)** Percent of fission products versus equivalent surface coverage for acetylated and unmodified αSyn-NTD. Error bars in panels A, C, E-F represent the standard deviation of the mean from 3 independent replicate experiments. Error bars in B,D represent the standard error of the mean.

To scale the K_*D,app*_, the lipid concentration in the imaging wells and in both SUPER template assays must be calculated, and then the ratios of these concentrations are used to scale the K_*D,app*_. The 2D concentration of lipids in the tethered vesicle assay was determined described below.

The total number of vesicles in each imaging well is given by the following equation:

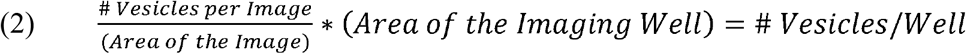

Where the area of each image was 1285 µm^2^.

The moles of lipid in each vesicle are given by the following equation:

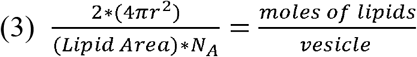

Where the lipid area is the average projected area of each lipid on the surface of the bilayer. This value was determined to be a weighted average between 63% DOPC (0.72 nm^2 38^), 25% DOPA, and 10% DGS-NTA. Although no reported lipid area exists for DOPA, its identical acyl chains and slightly smaller headgroup relative to DOPC suggest an estimated area of ~0.70 nm^2^. Similarly, the size of DGS-NTA can be estimated to be near DOPC at ~0.72 nm^2^. This results in an average area per lipid of 0.71 nm^2^. The average radius (r) of each vesicle was 104 nm, as determined by DLS. The factor of 2 on the left-hand side accounts for two leaflets in the bilayer, and N_A_ is Avogadro’s number. Multiplying Eqs. 2 and 3 yields the total moles of lipid per well. Dividing this product by the volume of the imaging well (30 µL) yields a lipid concentration of 60.0 nM.

A similar calculation was performed for the 9.2 µm SUPER templates in the imaging wells. Equation 1 was modified to account for the average number of SUPER templates per image with an image area of 20,995 µm. Equation 3 was modified using the template radii, and the final estimated lipid concentration in each well was 131.6 nM.

The final lipid concentration was calculated for the SUPER templates used for spectroscopy. 5 × 10^6^ templates were synthesized in 100 µL of solution. 5 µL of this template solution was used per condition in spectroscopy, resulting in 2.5 × 10^5^ beads per tube. Using Eq. 3, 1.25 µm radius for 2.5 µm beads, multiplying the result by the number of beads per tube and dividing by the solution volume (50 µL), yields a lipid concentration of 460 nM per tube.

The final lipid ratios between the tethered vesicle assay and each SUPER template assay were found by dividing the SUPER template concentrations by the tethered vesicle assay concentration for a ratio of 7.7 and 2.2 for the spectroscopy and microscopy assays, respectively. Bound protein density in the SUPER template assays was determined by rescaling the fitted K_*D,app*_ values using the lipid concentration ratios and substituting the adjusted K_*D,app*_ into Eq. 1 to calculate the number of bound proteins. This value was converted to surface density (proteins per 1000 nm^2^) by normalizing to the surface area of a 104 nm diameter vesicle.

### Quantification of Membrane Coverage and Thermodynamic Modeling of Membrane Fission

The radius of gyration (Rc) of αSyn-CTD and αSyn-CTDx3 were estimated using Eq. 4, which has previously been applied to and validated for a wide range of IDPs^39^. Within this equation, the IDP is treated as a self-avoiding chain in good solvent where ν = 0.588 and α= 0.2. N is equal to 43 and 139 for αSyn-CTD and αSyn-CTDx3, respectively (Supplementary Table S2).

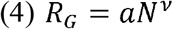

The projected membrane area of each membrane-bound construct was estimated from its radius of gyration according to Eq. 5. The calculated R_*G*_ and projected area for both αSyn-CTD constructs are shown in Supplementary Table S2.

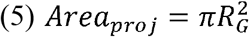

The projected area was then used to convert experimentally measured bound protein densities to fractional membrane coverage (η) using Eq. 6. This is a dimensionless parameter that represents the fraction of membrane area occupied by the intrinsically disordered region.

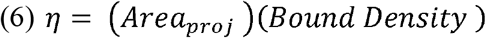

Membrane fission was modeled using a coverage-dependent free energy described in Eq. 7 where ΔG_*o*_ corresponds to the baseline level of membrane fission occurring in the absence of bound protein and *A* is a regressed parameter that describes the magnitude of the IDP-mediated crowding pressure. The 9/4 scaling arises from, and 9/4 arises from Des Cloiseaux scaling laws for polymers ^40–41^ (ref) and has previously been validated for membrane-bound IDPs^42^.

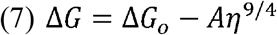

The fission percentage was estimated to follow a Boltzmann distribution as shown in Eq. 8 using the ΔG calculated in Eq. 7 with ka (Boltzmann’s constant) and T (absolute temperature).

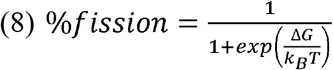

Nonlinear least-squares regression was used to determine the optimal value of ΔG_*o*_ and *A* for each experimental condition. For both salt conditions, ΔG_*o*_ was constrained to be identical for αSyn-CTD and αSyn-CTD×3 because of identical membrane templates, but each salt condition was allowed distinct ΔG_*o*_ due to the vast difference in salt concentration (4.59 k_*B*_ *T* for 150 mM NaCl and 4.53 k_*B*_ *T* for 1000 mM NaCl). *A* was allowed to vary between protein constructs to determine if the curves would collapse onto a coverage-dependent curve. For 150 mM NaCl, A = 1027 k_*B*_ *T* for αSyn-CTD and 171 k_*B*_ *T* for αSyn-CTDx3. For 1000 mM NaCl, A = 99 k_*B*_ *T* for both constructs.

## RESULTS AND DISCUSSION

To dissect the contributions of individual domains to membrane fission and remodeling, we generated truncation mutants from full-length αSyn (αSyn-FL, 140 residues) (Fig. 1B). The first 97 amino acids were used to create αSyn-NTD and residues 98-140 were used to synthesize αSyn-CTD. Throughout this study, we utilized the N-terminally acetylated αSyn-FL and αSyn-NTD, as this post-translational modification represents the physiologically relevant form of αSyn *in vivo* ^43–45^. An N-terminal hexa-histidine tag was appended to αSyn-CTD to enable membrane recruitment through DGS-NTA lipids (Figs. 1B-C), as the CTD lacks intrinsic membrane binding motifs. All protein amino acid sequences can be found in Supplementary Table 1. Lipid vesicles were composed primarily of DOPC with DOPA (25 mol%) and DGS-NTA (10 mol%) included to elicit binding from all protein constructs.

SUPER templates were utilized to analyze membrane remodeling behavior of each αSyn construct (Fig. 1D). The inclusion of ATTO647N-DPPE in membranes (1 mol%) allowed for high throughput quantification of membrane release via a fluorescence plate reader (see methods) ^28, 33^, as well as the visualization of protein-induced membrane morphological changes via fluorescence microscopy ^33^.

Membrane fission was quantified in Figure 2. All protein constructs elicited monotonic increases in ATTO647N supernatant lipid fluorescence with increasing protein concentration, consistent with fission-induced membrane release (Fig. 2A, see Methods). αSyn-FL exhibited the highest degree of membrane fission, with apparent saturation above 1000 nM protein concentration. The lesser degrees of membrane fission induced by αSyn-NTD and αSyn-CTD indicate that both domains contribute to αSyn-FL-mediated fission simultaneously. Since membrane remodeling behavior is known to be a function of protein binding density ^17, 46^, we controlled for the differing levels of membrane affinity by generating binding curves for each protein construct (Fig. 2B). αSyn-CTD, which relies on high affinity hexahistidine-Ni^2+^ interactions for binding^47–48^, had the highest membrane affinity of the three protein constructs. αSyn-NTD likely had stronger affinity for the membrane than αSyn-FL due to its absence of the highly anionic CTD, which would Coulombically oppose binding to the anionic, DOPA-rich membranes used here. When membrane affinity was accounted for in Figure 2C, the trend of αSyn-FL eliciting the highest degree of membrane fission was maintained. This result indicates that the NTD and CTD of αSyn possess distinct, intrinsic abilities to drive membrane fission.

Membrane fission by αSyn-CTD likely arises from crowding-related steric and/or electrostatic effects that have been previously demonstrated for membrane-tethered IDPs ^22–24^. For αSyn-NTD, the ability to induce fission may result from a combination of steric crowding and asymmetric membrane insertion by the amphipathic helix ^22, 25, 28, 46^. To further probe the impact of amphipathic helical insertion on membrane fission, we tested αSyn-FL in both the acetylated and unmodified forms. The presence of N-terminal acetylation marginally enhanced affinity on DOPA-containing membranes (Fig. 2D), consistent with previously reported effects of this modification on moderately charged membranes ^32, 49^. However, no appreciable differences in fission efficacy were observed for the acetylated and unmodified forms of αSyn-FL (Fig. 2E, Fig S2A). Similar results were observed for αSyn-NTD (Fig. 2F, Fig. S2B-C). Together, these results indicate that although N-terminal acetylation modestly alters membrane binding, it has little impact on fission efficacy, suggesting that αSyn-mediated membrane remodeling is governed primarily by domain architecture of the NTD and CTD.

While the fluorescence-based SUPER template assay provides a quantitative measure of membrane fission, it does not report on other morphological changes associated with membrane remodeling. We therefore used fluorescence microscopy to directly visualize protein-induced membrane curvature on the surface of templates with diameters of approximately 10 μm (Fig. 3). Figure 3A shows single representative confocal imaging slices of the lipid, protein, and merged fluorescent channels for templates in the absence or presence of a protein construct. By analyzing the z-stack images in each condition, we quantified the percentage of templates with optically resolvable surface features, such as tubulation and vesiculation (Figs. 3B-C). Representative movies capturing the highly dynamic nature of these surface features are provided in the supporting information (Movie S1).

**Figure 3.**
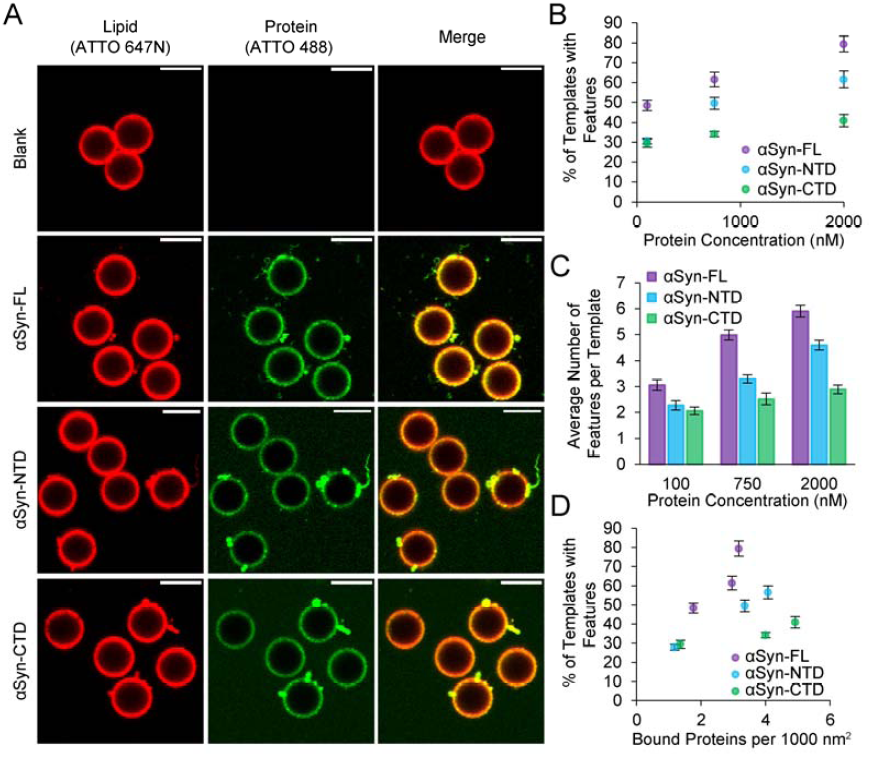
Quantification of membrane morphological deformation by αSyn truncation mutants. **(A)** Representative confocal fluorescence micrographs of SUPER templates incubated with 2 µM protein. Scale bars = 10 µm. **(B)** Percentage of templates exhibiting membrane features as a function of protein concentration. **(C)** Average number of features per remodeled template as a function of protein concentration. **(D)** Percentage of templates exhibiting membrane features as a function of bound protein density. Error bars represent the standard error of the mean (N = 144-323 imaged templates per condition).

Consistent with the results from Figures 2A-C, Figure 3B shows that the percentage of imaged templates displaying membrane features increases with increasing protein concentration for all αSyn constructs. As observed in the fission assays, αSyn-FL exhibits the strongest remodeling behavior, while αSyn-CTD induces the least membrane deformation. A qualitatively similar trend was observed when counting the average number of curved features per template (Fig. 3D). When differences in membrane binding between the constructs are accounted for, these trends persist (Fig. 3C), further indicating that variations in membrane affinity are not responsible for the observed remodeling behavior. Similar to the results from Figures 2D-F, N-terminal acetylation was shown to have negligible impact on the degree of membrane deformation observed (Fig. S3). These results reiterate the synergy between αSyn-NTD and αSyn-CTD that gives rise to the full remodeling ability of αSyn-FL. Notably, templates in the absence of protein tended to cluster (Fig. 3A), whereas protein-treated templates remained spatially dispersed, particularly in the αSyn-FL and αSyn-CTD conditions, consistent with CTD-mediated steric stabilization of membrane surfaces^50^.

To better understand the energetics underlying αSyn-mediated membrane remodeling, we next classified the types of membrane structures observed on the template surfaces. The membrane features observed on the template surfaces fell into three main morphological categories: membrane buds, thick tubules, and sub-diffraction tubules (Fig. 4A). For clarity, the representative image slices highlight a single feature per template, although individual templates typically exhibit multiple features, as quantified in Fig. 3D. Here, membrane buds refer to approximately spherical protrusions extending from the membrane surface, while thick tubules correspond to cylindrical membrane structures with diameters that are optically resolvable by confocal microscopy. In contrast, sub-diffraction tubules refer to highly curved tubular structures whose diameters fall below the optical diffraction limit and therefore appear as diffraction-limited features in the fluorescence images.

**Figure 4.**
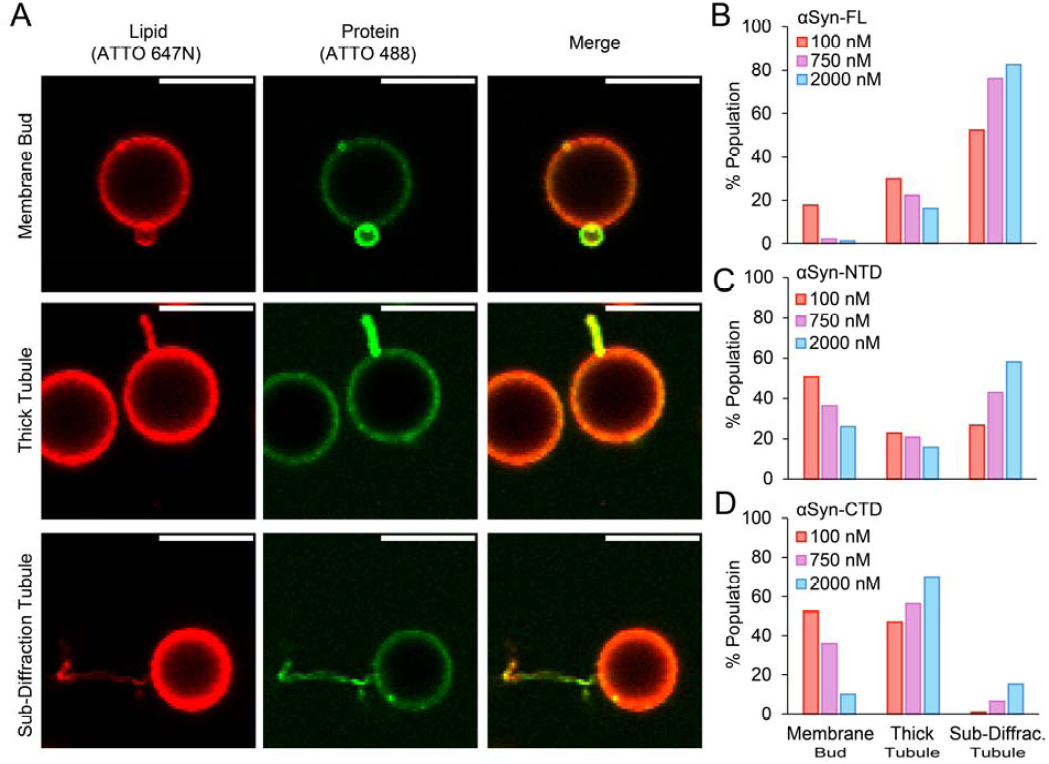
Distribution of membrane morphological features induced by αSyn truncation mutant binding. **(A)** Representative confocal fluorescence micrographs of membrane buds, thick tubules, and sub-diffraction tubules formed on SUPER templates. Scale bars = 10 µm. Distribution of feature types as a function of protein concentration for **(B)** αSyn-FL, **(C)** αSyn-NTD, and **(D)** αSyn-CTD.

These morphological classes correspond to progressively higher curvature deformations of the membrane. Because membrane bending energy increases with curvature, sub-diffraction tubules represent higher-energy deformations than thick tubules or membrane buds. Within the Helfrich framework, bending energy scales with the square of curvature, implying that narrower tubules require greater energetic input ^51–52^. Accordingly, as the concentration of αSyn-FL increases, the distribution of observed structures shifts towards progressively higher-curvature morphologies (i.e., sub-diffraction tubules > thick tubules > membrane buds) (Fig. 4B). Specifically, both membrane buds and thick tubules become depleted while sub-diffraction tubules become the dominant feature at the highest protein concentration. A qualitatively similar trend is observed for αSyn-NTD (Fig. 4C), although the shift toward higher-curvature morphologies is less pronounced. At lower protein concentrations, a larger fraction of the observed structures are membrane buds than in the corresponding conditions with αSyn-FL in Figure 4B. A similar distribution of membrane buds was observed for templates incubated with αSyn-CTD (Fig. 4D). However, increasing αSyn-CTD concentration primarily leads to the formation of thick tubules, which becomes the dominant morphology at higher protein concentrations. Although the proportion of sub-diffraction tubules also increases with αSyn-CTD concentration, they represented only a small fraction of the overall structure population

Taken together, these distributions highlight clear differences in the remodeling behavior of the three αSyn constructs. αSyn-FL shows the strongest shift toward sub-diffraction tubules, whereas αSyn-CTD remains enriched in lower-curvature structures, primarily membrane buds and thick tubules. αSyn-NTD exhibits an intermediate distribution, with a more balanced presence of all three membrane structure types. This trend suggests that the amphipathic helix of the NTD plays an important role in driving membranes toward highly curved morphologies, as observed for both αSyn-FL and αSyn-NTD. At the same time, the additional presence of the CTD in the full-length protein further enhances the shift toward the highest-curvature structures. These results are consistent with the idea that the two domains of αSyn act cooperatively to amplify the overall membrane remodeling ability of the protein. When these membrane structure categories were further subdivided based on tubule length, yielding five morphological categories in total, the resulting distributions qualitatively followed the same trends observed above (Figure S4). Notably, protrusive membrane features often exhibited elevated protein fluorescence relative to the surrounding template surface (Fig. 4A). While this result may suggest enrichment of αSyn on curved structures, curved membranes can also appear brighter simply because their geometry places more fluorophores within the optical path.

While the remodeling activity of αSyn-NTD can be readily attributed to amphipathic helix insertion, the mechanism by which αSyn-CTD drives membrane deformation remains less clear, particularly given that it is a short and highly charged domain lacking intrinsic membrane insertion motifs. To investigate this behavior, we examined two potential contributions to αSyn-CTD-mediated remodeling: steric repulsion and electrostatic repulsion (Fig. 5). To probe the steric contributions of the CTD, we generated a construct containing three tandem repeats of the CTD sequence fused (αSyn-CTDx3), thereby increasing the effective length of the disordered domain while maintaining the same charge density. The Y136C substitution (numbering relative to αSyn-FL) was introduced only in the final repeat to maintain consistency with the mutation used in αSyn-CTD (Supplementary Table 1). Increasing the length of the disordered domain substantially enhanced membrane remodeling. Across comparable membrane binding densities, αSyn-CTDx3 produced greater membrane fission than αSyn-CTD (Fig. 5A, Fig S5A-B). Similarly, αSyn-CTDx3 increased both the fraction of templates exhibiting membrane features and the average number of features per template (Figs. 5B-5C, Fig S5C). These results demonstrate that increasing the length of the disordered CTD amplifies its ability to drive membrane deformation.

**Figure 5.**
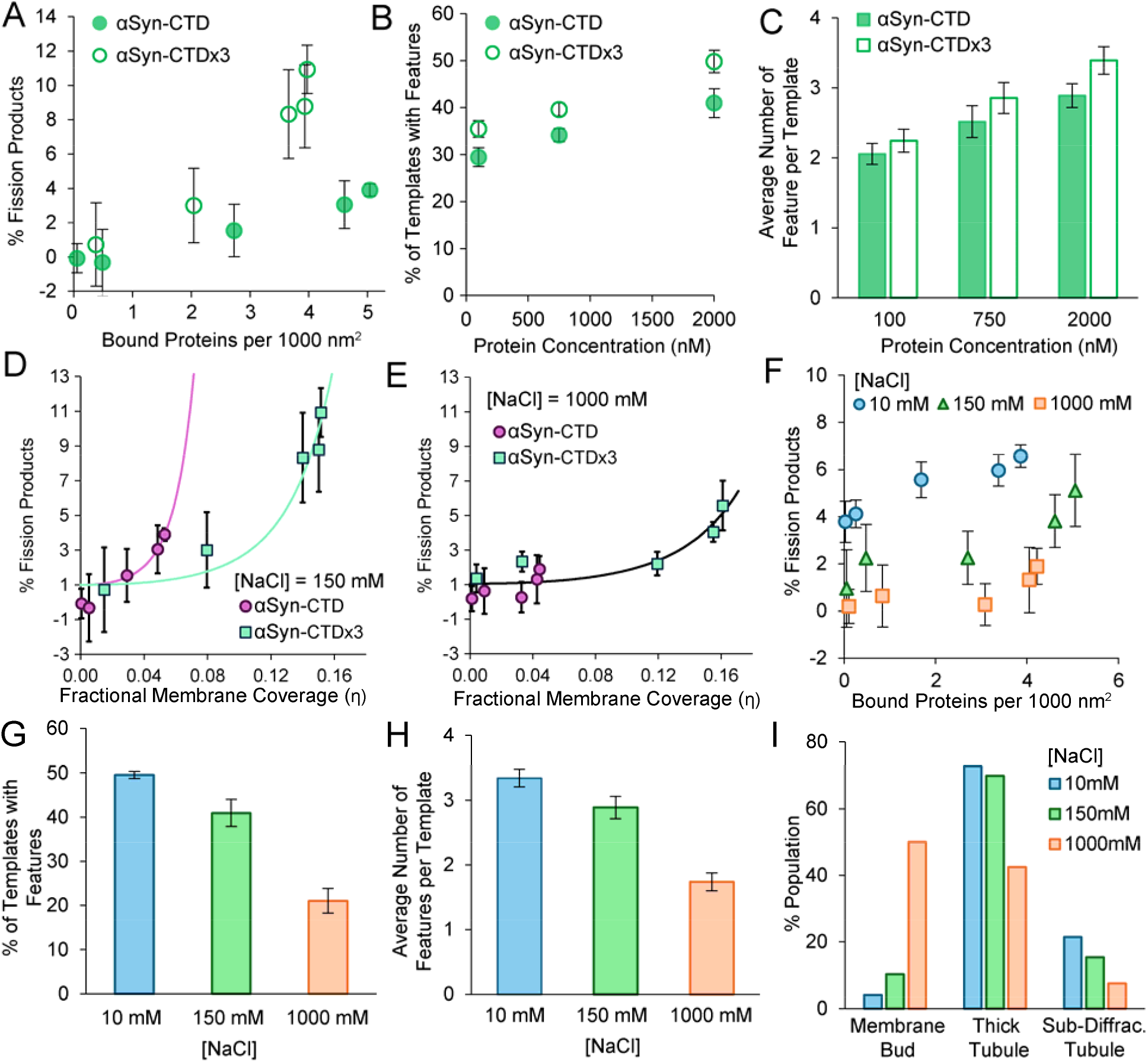
Steric and electrostatic contributions of αSyn-CTD to membrane remodeling. **(A)** Percent of fission products generated by either αSyn-CTD and αSyn-CTDx3 as a function of bound protein density. **(B)** Percentage of templates exhibiting membrane features as a function of protein concentration. **(C)** Average number of features per remodeled template as a function of protein concentration. **(D)** Percent of fission products as a function of membrane coverage for both αSyn-CTD and αSyn-CTDx3 at 150 mM NaCl. **(E)** Percent of fission products as a function of membrane coverage for both αSyn-CTD and αSyn-CTDx3 at 1000 mM NaCl. **(F)** Percent of fission products plotted as a function of bound protein density for different ionic strengths. **(G)** Percentage of templates exhibiting membrane features as a function of salt concentration in the presence of 2 µM αSyn-CTD. **(H)** Average number of features per remodeled template as a function of salt concentration in the presence of 2 µM αSyn-CTD. **(I)** The distribution of feature types on all templates as a function of salt concentration in the presence of 2 µM αSyn-CTD. Error bars represent standard deviation of the mean from 3 independent replicate measurements. Error bars in A-C and G-H represent the standard error of the mean (N = 190-323 imaged templates).

To determine whether this enhancement arises purely from steric crowding or disordered chains on the membrane surface, we analyzed membrane fission as a function of fractional membrane coverage (η). Here η is simply the product of the bound protein surface density (i.e., proteins per nm^2^) and the projected area of a single protein, which was estimated from established scaling laws^39^ for soluble IDPs using chain lengths of 43 and 129 residues for αSyn-CTD and αSyn-CTDx3, respectively (see Methods). Previous work applying polymer brush theory to membrane-tethered IDPs^42^ demonstrated that steric pressure in the semi-dilute regime follows the Des Cloiseaux scaling law^40–41^, in which the free energy scales with η^9/4^ and becomes largely independent of chain length. Consistent with this framework, IDPs of different lengths collapse onto a single scaling curve when steric interactions dominate^42^.

To test whether αSyn-CTD remodeling follows this behavior, we converted the predicted free-energy dependence into an expected fraction of fission products using a Boltzmann distribution and examined whether αSyn-CTD and αSyn-CTDx3 collapse onto a common scaling relationship (Figs. 5D-E). At physiological ionic strength (150 mM NaCl), the two constructs fail to collapse onto a single curve, with αSyn-CTD exhibiting stronger dependence on membrane coverage than αSyn-CTDx3 (Fig. 5D). This deviation from the expected scaling behavior indicates that steric crowding alone cannot fully account for the observed remodeling activity. In contrast, when electrostatic interactions were screened by increasing the salt concentration to 1 M NaCl, the data from αSyn-CTD and αSyn-CTDx3 collapsed onto a common scaling curve (Fig. 5E), consistent with the predictions of steric polymer crowding. The lack of collapse under physiological ionic strength in Figure 5D therefore suggests that electrostatic interactions between neighboring IDPs substantially contribute to membrane remodeling.

To probe electrostatic contributions further, we systematically varied the ionic strength of the solution and quantified the resulting effects of membrane remodeling by αSyn-CTD. Membrane fission was strongly modulated by salt concentration, with substantially reduced fission observed at 1000 mM NaCl and enhanced fission at 10 mM NaCl relative to physiological conditions (150 mM NaCl) (Fig. 5F, Fig. S5D-E). Consistent with these fission measurements, membrane morphology also exhibited a pronounced dependence on ionic strength. At constant αSyn-CTD concentration, decreasing ionic strength increased both the fraction of templates displaying deformed membrane features (Fig. 5G) and the average number of features per template (Fig. 5H). Increasing ionic strength yielded the opposite effect, suppressing the formation of curved structures. Qualitatively similar results were observed for αSyn-CTDx3 (Figure S6A-D).

Changes in ionic strength also altered the distribution of membrane morphologies (Fig. 5I, Fig. S6E). Lower salt concentrations favored high-curvature structures, while strong electrostatic screening shifted the population toward lower-energy morphologies. Together, these results demonstrate that CTD-mediated membrane remodeling is highly sensitive to ionic strength, indicating that electrostatic repulsion between membrane-tethered CTD domains plays a significant role in driving membrane deformation.

Taken together, the results in Figures 2–5 define a mechanistic picture of αSyn-mediated membrane remodeling. The N-terminal amphipathic helix generates an initial curvature bias through asymmetric membrane insertion, while the membrane-tethered CTD produces lateral pressure that arises primarily from electrostatic repulsion between neighboring, highly charged disordered domains. In the full-length protein, these mechanisms work in concert to cooperatively enhance membrane remodeling. Although the NTD appears to be the stronger curvature-inducing element, the CTD supplies a non-negligible energetic contribution that enhances remodeling efficiency. This cooperation suggests that post-translational modifications or mutations that alter CTD charge density, such as phosphorylation or truncation ^53–54^, could modulate curvature generation by enhancing or weakening electrostatic pressure on the membrane surface.

## CONCLUSION

In this work, we show that membrane remodeling by α-synuclein emerges from a cooperative interplay between its structured NTD and intrinsically disordered CTD. The NTD drives curvature through amphipathic helix insertion, establishing an initial membrane bias. By modulating ionic strength, changing the IDP chain length, and incorporating polymer scaling analysis, we determined that the membrane-tethered CTD generates lateral pressure through electrostatic protein-protein repulsion, further amplifying membrane deformation. Together, these domains produce greater fission and higher-curvature morphologies than either domain alone. More broadly, these findings refine physical models of intrinsically disordered protein– membrane interactions by demonstrating that relatively short, disordered domains can exert substantial bending stresses when densely charged and membrane-bound. The CTD, therefore, functions as a tunable energetic amplifier of membrane curvature, suggesting that sequence modifications that alter charge density may directly regulate αSyn-mediated remodeling in physiological and pathological contexts.

## ASSOCIATED CONTENT

### Supporting Information

The following files are available free of charge.

Additional experimental data, protein construct sequences, and thermodynamic parameters for membrane fission modeling (PDF)

Movie of the dynamic protrusions on SUPER template surfaces (AVI)

## AUTHOR INFORMATION

### Author Contributions

D.J. and W.Z. designed experiments, D.J. and J.L. performed experiments, and all authors contributed to manuscript preparation.

## Notes

The authors declare no competing interests.

## ACKNOWLEDGMENTS

Thank you to Professor Peter Chung and Antonis Margaritakis at the University of Southern California for allowing us to use their plate reader to perform this work. This research was supported by the National Institutes of Health through R35GM147333 to D.J., O.K., and W.Z.

