## Supporting Information for "The disordered and structured regions of α-Synuclein contribute to membrane remodeling synergistically"

*Corresponding Author: Wade F. Zeno

| | |
|---|---|
| **αSyn-FL (wildtype)** | MDVFMKGLSKAKEGVVAAAEKTKQGVAEAAGKTKEGVLYVGSKTKEGV<br>VHGVATVAEKTKEQVTNVGGAVVTGVTAVAQKTVEGAGSIAAATGFVKK<br>DQLGKNEEGAPQEGILEDMPVDPDNEAYEMPSEEGYQDYEPEA |
| **αSyn-FL (Y136C mutant)** | MDVFMKGLSKAKEGVVAAAEKTKQGVAEAAGKTKEGVLYVGSKTKEGV<br>VHGVATVAEKTKEQVTNVGGAVVTGVTAVAQKTVEGAGSIAAATGFVKK<br>DQLGKNEEGAPQEGILEDMPVDPDNEAYEMPSEEGYQDCEPEA |
| **αSyn-NTD** | MDVFMKGLSKAKEGVVAAAEKTKQGVAEAAGKTKEGVLYVGSKTKEGV<br>VHGVATVAEKTKEQVTNVGGAVVTGVTAVAQKTVEGAGSIAAATGFVKK |
| **αSyn-NTD (G93C mutant)** | MDVFMKGLSKAKEGVVAAAEKTKQGVAEAAGKTKEGVLYVGSKTKEGV<br>VHGVATVAEKTKEQVTNVGGAVVTGVTAVAQKTVEGAGSIAAATCFVKK |
| **αSyn-CTD (Y136C mutant)** | GSHHHHHHGSGSDQLGKNEEGAPQEGILEDMPVDPDNEAYEMPSEEG<br>YQDCEPEA |
| **αSyn-CTDx3 (Y136C mutant)** | GSHHHHHHGSGSDQLGKNEEGAPQEGILEDMPVDPDNEAYEMPSEEG<br>YQDYEPEADQLGKNEEGAPQEGILEDMPVDPDNEAYEMPSEEGYQDYE<br>PEADQLGKNEEGAPQEGILEDMPVDPDNEAYEMPSEEGYQDCEPEA |

**Supplementary Table 1.** αSyn construct amino acid sequences. The locations of cysteine point mutations are highlighted in yellow.

| | Charge Density (charge/length) | IDP Length | Radius of Gyration ($R_G$) (nm) | Projected Area ($nm^2$/protein) |
|---|---|---|---|---|
| **αSyn-CTD** | -0.32 | 43 | 1.82 | 10.47 |
| **αSyn-CTDx3** | -0.32 | 129 | 3.48 | 38.13 |

**Supplementary Table 2.** Calculated physical parameters of the αSyn-CTD constructs.

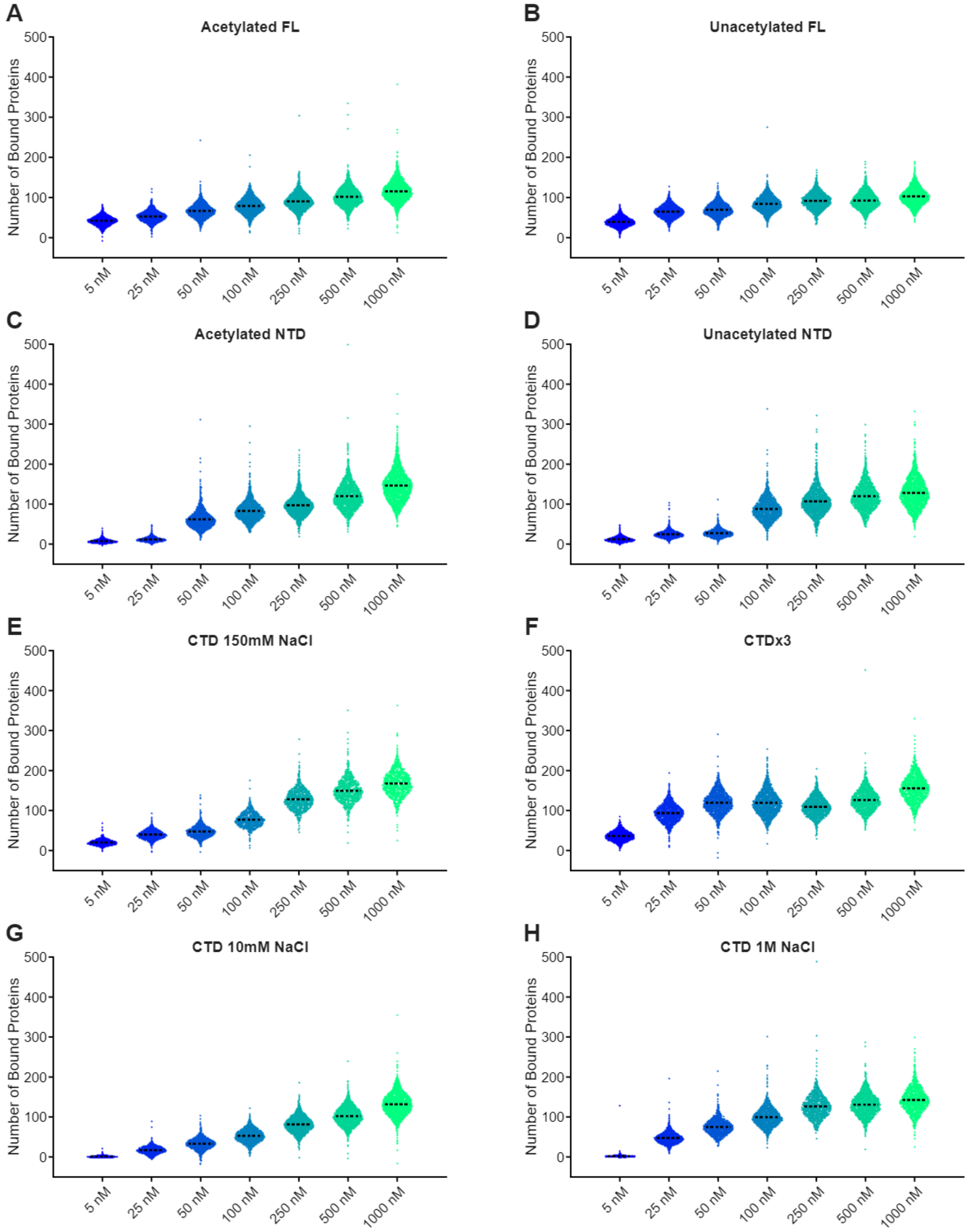


**Supplementary Figure 1**. Distributions of proteins bound to vesicles with average diameters of 100 nm. These distributions are the raw data for plots generated in Figures

2, 5, S2, and S5 of the main text. The black dashed lines indicate the average number of proteins bound.

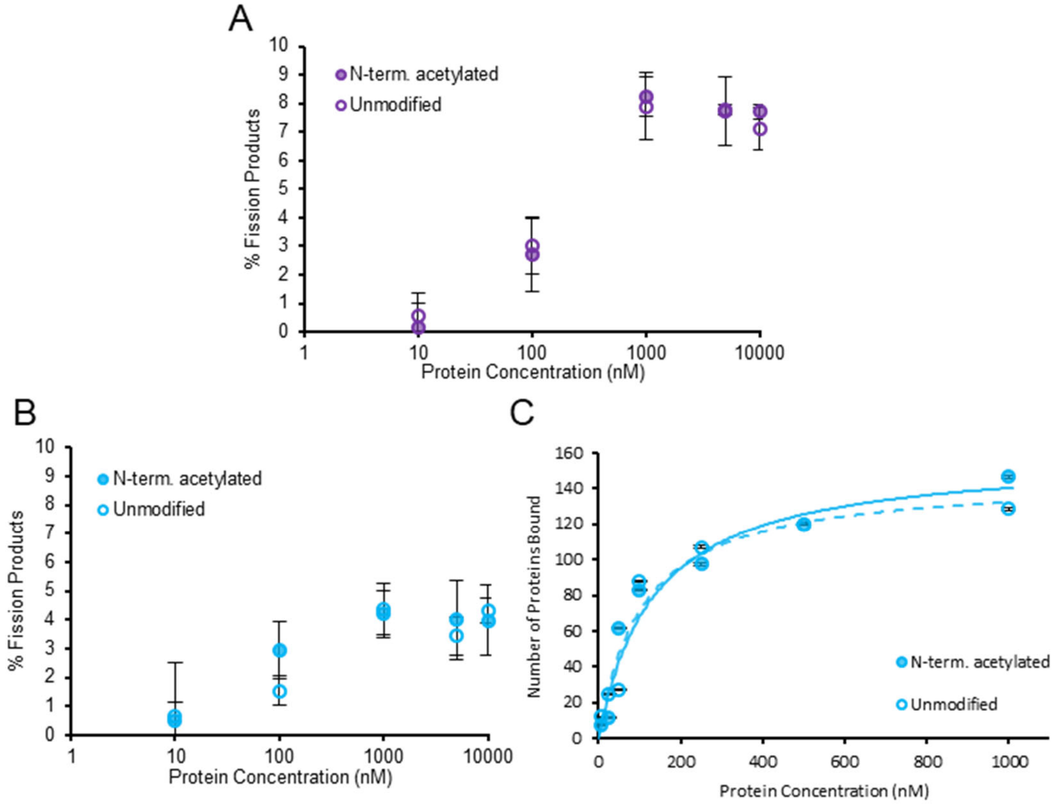


**Supplemental Figure 2. (A)** Percent of fission products for acetylated and unmodified αSyn-FL as a function of protein concentration. **(B)** Percent of fission products for acetylated and unacetylated αSyn-NTD as a function of concentration. **(C)** Binding isotherms obtained from tethered vesicle assays with Langmuir fits. N = 1579 – 2117 imaged vesicles for acetylated αSyn-NTD and N = 1631 – 2044 imaged vesicles for unmodified αSyn-NTD for each point on the binding curve. Error bars in A-B represent the standard deviation of the mean from 3 independent replicate experiments. Error bars in C represent the standard error of the mean (N = 1579 - 2093 imaged vesicles for each experimental condition).

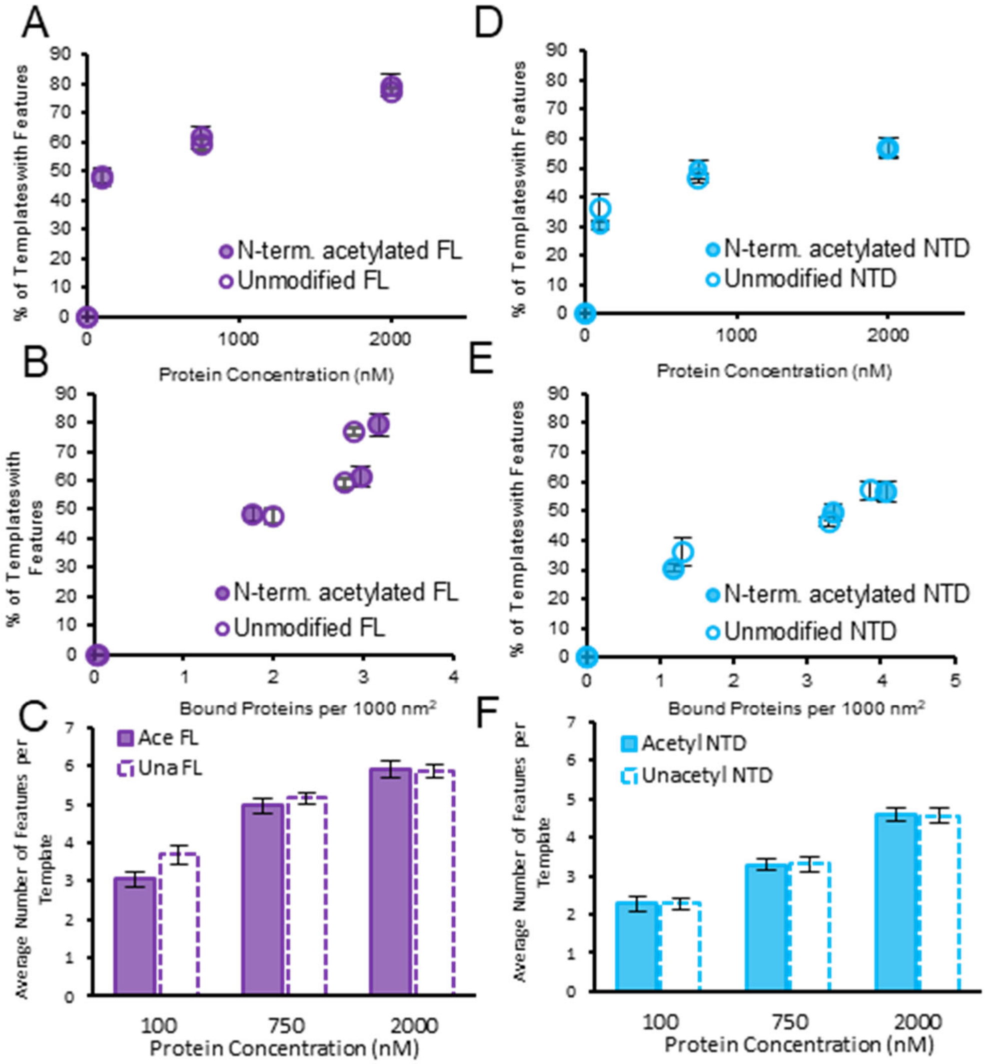


**Supplemental Figure 3**. Quantification of membrane morphological deformation induced by **(A-C)** acetylated αSyn constructs and **(D-F)** unmodified constructs. Error bars represent the standard deviations of the mean from N = 144 – 305 imaged templates for each experimental condition.

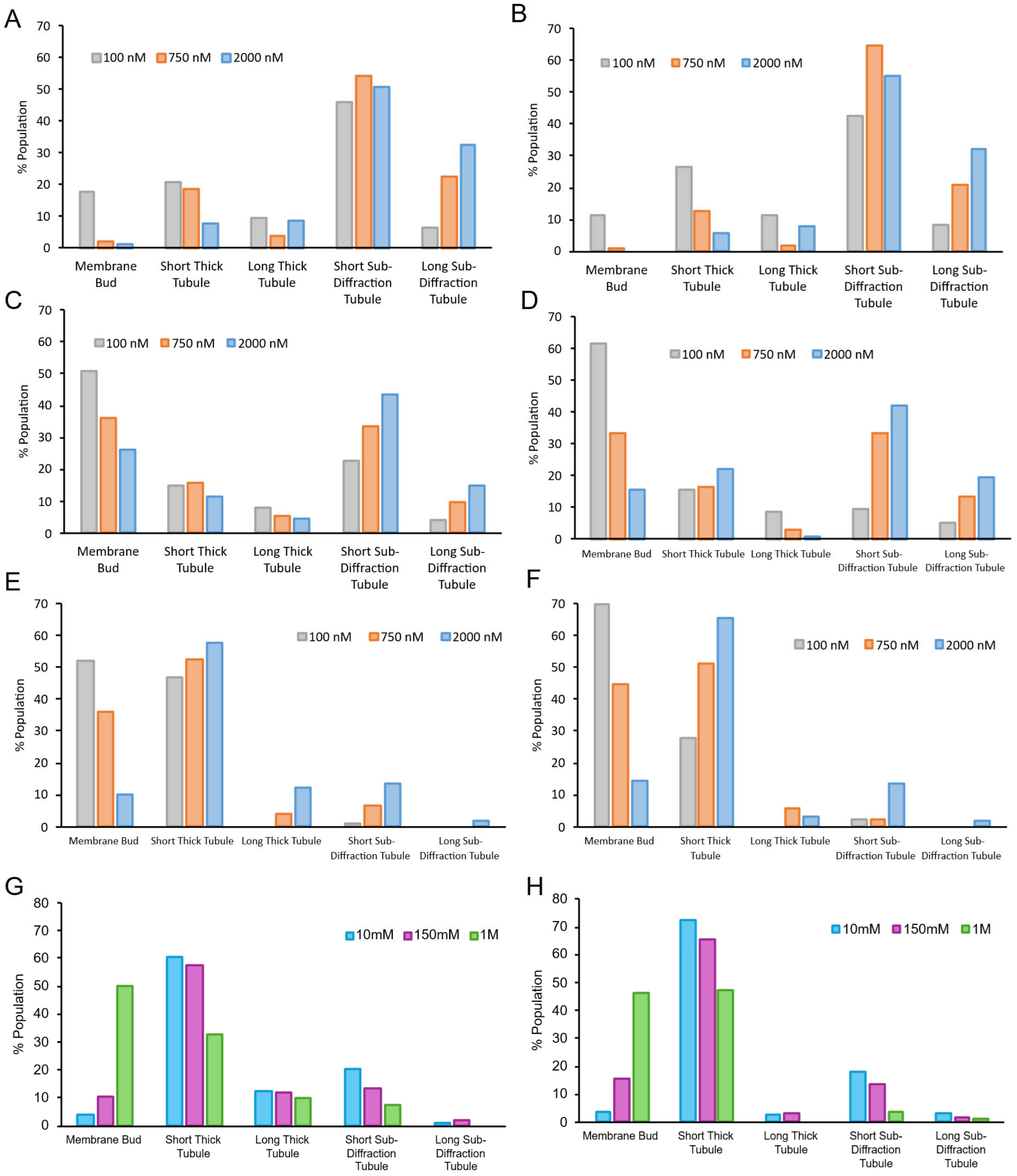


**Supplemental Figure 4.** Population distributions of features on SUPER templates. **(A)** Acetylated αSyn-FL **(B)** Unmodified αSyn-FL **(C)** Acetylated αSyn-NTD **(D)** Unmodified αSyn-NTD **(E)** αSyn-CTD **(F)** αSyn-CTDx3 **(G)** αSyn-CTD **(H)** αSyn-CTDx3. All proteins were incubated at a concentration of 2 μM.

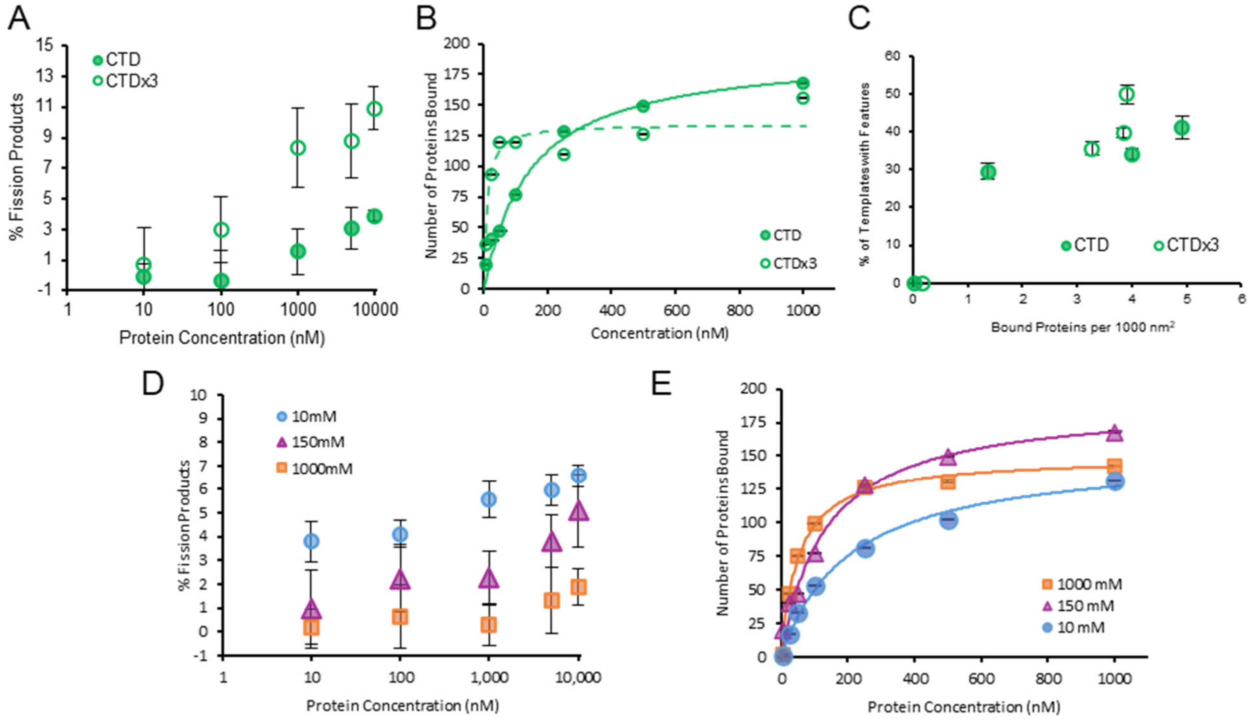


**Supplemental Figure 5. (A)** Percentage of membrane fission products as a function of protein concentration for αSyn-CTD and αSyn-CTDx3. **(B)** Binding isotherms with Langmuir fits obtained from tethered vesicle assay. N = 886-996 imaged vesicles for αSyn-CTD and N = 1351 – 1613 imaged vesicles for αSyn-CTDx3. **(C)** The data in Figure 5B plotted as a function of bound protein density. **(D)** Percent of fission products for αSyn-CTD as a function of protein concentration for various ionic strength in solution. **(E)** Binding isotherm for αSyn-CTD at varying ionic strengths. N = 1318 - 1917 imaged vesicles for 10mM NaCl, N = 886-996 imaged vesicles for 150mM NaCl, and N = 1069 – 1614 imaged vesicles for 1000 mM NaCl for each point on the isotherm. Error bars in A,D represent the standard deviation of the mean for 3 independent replicate experiments. The error bars in C represent the standard error of the mean for N = 190-323 imaged templates in each experimental condition.

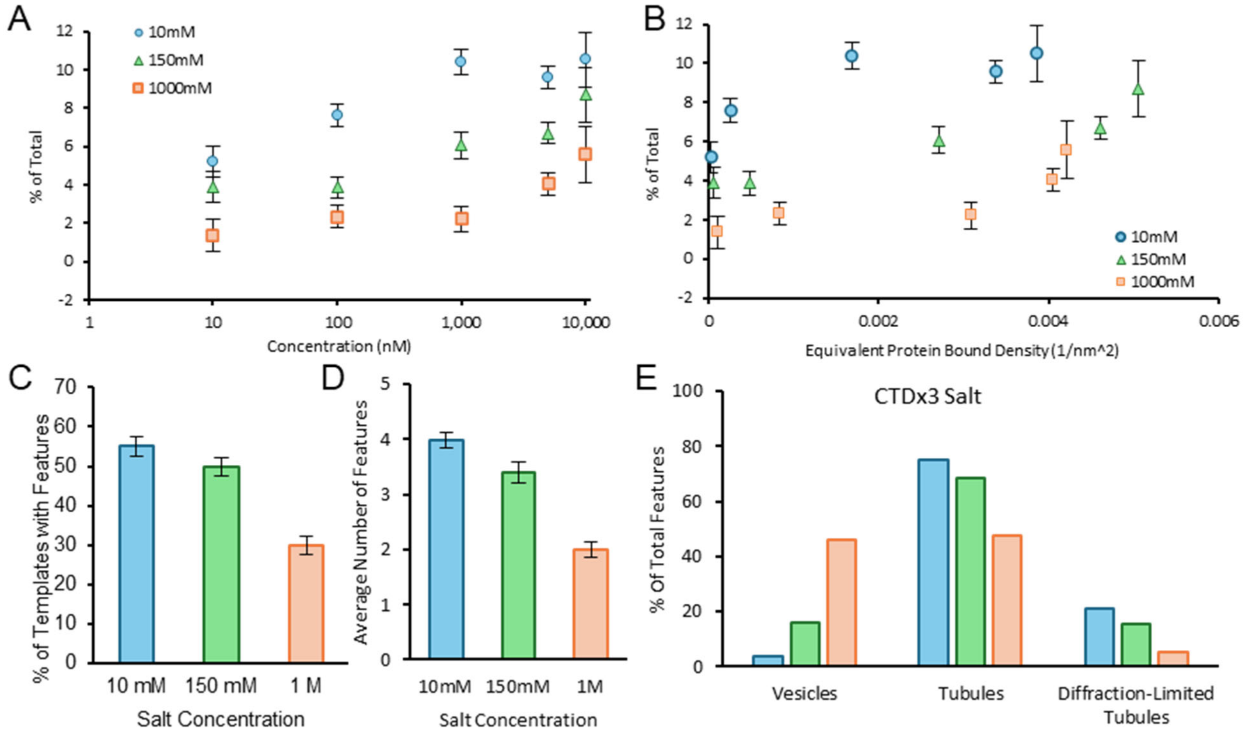


**Supplemental Figure 6**. Quantification of membrane remodeling for αSyn-CTDx3. **(A)** Percent membrane fission versus protein concentration under different ionic conditions. **(B)** The same data as (A) plotted as a function of bound protein density. **(C)** Percentage of templates exhibiting membrane features as a function of ionic strength. **(D)** Average number of deformation features per template as a function of salt concentration**. (E)** The distribution of feature type on all templates as a function of salt concentration. Error bars in A,B represent the standard deviation of the mean for 3 independent replicate experiments. Error bars in C-D represent the standard error of the mean for N = 120-267 imaged templates for each experimental condition.
